# Reliability of transcranial magnetic stimulation evoked potentials to detect the effects of theta-burst stimulation of the prefrontal cortex

**DOI:** 10.1101/2021.12.11.472198

**Authors:** Adriano H. Moffa, Stevan Nikolin, Donel Martin, Colleen Loo, Tjeerd W. Boonstra

## Abstract

**Background:** Transcranial magnetic stimulation (TMS) with simultaneous electroencephalography (EEG) is a novel method for assessing cortical properties outside the motor region. Theta burst stimulation (TBS), a form of repetitive TMS, can non-invasively modulate cortical excitability and has been increasingly used to treat psychiatric disorders by targetting the dorsolateral prefrontal cortex (DLPFC). The TMS-evoked potentials (TEPs) analysis has been used to evaluate cortical excitability changes after TBS. However, it remains unclear whether TEPs can detect the neuromodulatory effects of TBS.

**Objectives:** To confirm the reliability of TEP components within and between sessions and to measure changes in neural excitability induced by intermittent (iTBS) and continuous TBS (cTBS) applied to the left DLPFC.

**Methods:** Test-retest reliability of TEPs and TBS-induced changes in cortical excitability were assessed in twenty-four healthy participants by stimulating the DLPFC in five separate sessions, once with sham and twice with iTBS and cTBS. EEG responses were recorded of 100 single TMS pulses before and after TBS, and the reproducibility measures were quantified with the concordance correlation coefficient (CCC).

**Results:** The N100 and P200 components presented substantial reliability within the baseline block (CCCs>0.8) and moderate concordance between sessions (CCC_max_≈0.7). Both N40 and P60 TEP amplitudes showed little concordance between sessions. Changes in TEP amplitudes after iTBS were marginally reliable for N100 (CCC_max_=0.52), P200 (CCC_max_=0.47) and P60 (CCC_max_=0.40), presenting only fair levels of concordance at specific time points.

**Conclusions:** The present findings show that only the N100 and P200 components had good concordance between sessions. The reliability of earlier components may have been affected by TMS-evoked artefacts. The poor reliability to detect changes in neural excitability induced by TBS indicates that TEPs do not provide a precise estimate of the changes in excitability in the DLPFC or, alternatively, that TBS did not induce consistent changes in neural excitability.

## 1. Introduction

In recent years, TMS-evoked potentials (TEPs) have gained ground among researchers working with non-invasive brain stimulation (NIBS) techniques as a method to non-invasively assess cortical excitability and connectivity properties beyond motor regions (Tremblay et al., 2019). This approach uses a combination of electroencephalography (EEG) and single/paired pulses of transcranial magnetic stimulation (TMS) (Ilmoniemi et al., 1997). TEPs have been increasingly used to assess the brain state of healthy and pathological individuals, and evaluate and monitor the impact of pharmacological and NIBS interventions on brain activity and cortical circuits (Cao et al., 2021, Chung et al., 2015). Among NIBS techniques, theta-burst stimulation (TBS), a form of repetitive TMS, can modulate cortical excitability and has been increasingly used to treat psychiatric disorders involving the dorsolateral prefrontal cortex (DLPFC). However, neither the reliability of TEP measures nor TBS-induced effects on the excitability of the DLPFC have been fully elucidated. This represents a critical limitation for further experimental and clinical studies applying TBS to the DLPFC or using TEPs to investigate prefrontal cortical excitability. The present study aims to address these gaps in knowledge, estimating the test-retest reliability of the TMS-EEG methodology and TBS-induced changes in cortical excitability of the DLPFC.

To date, two studies have assessed the reliability of TEPs in the motor cortex. Lioumis et al. (2009) found a significant correlation between test-retest measurements with TMS-EEG responses achieving high overall reproducibility. Casarotto et al. (2010) evaluated the sensitivity and repeatability of concurrent TMS-EEG measures in 10 healthy volunteers showing that TEPs were sensitive to changes in stimulation parameters and repeatable over time. While these initial findings are promising, the restriction to the motor cortex, the small sample sizes, and the limitations of some of the reliability metrics used, mean the test-retest reliability of TEPs still requires further evaluation. For example, the degree of correlation between repeated TEP measures is necessary but may not be sufficient to confirm measurement agreement (Lawrence and Lin, 1989) and is therefore not considered an ideal measure of reliability (Koo and Li, 2016). More recently, Kerwin et al. (2018) assessed the reliability of TEPs resulting from stimulation of the left DLPFC using the concordance correlation coefficient (CCC), which assesses the ability of a test to distinguish between different individuals. CCC is specifically designed to assess reliability between repeated measures as it considers both accuracy and precision (Lawrence and Lin, 1989). The main findings were that TEPs were generally robust and reliable, with the best reliability levels achieved for the N100 and P200 components.

While these studies investigated the reliability of the TEP components themselves, they did not assess whether TEPs can reliably measure changes in cortical excitability. NIBS has been widely used to induce changes in cortical excitability. For example, theta-burst stimulation involves repeated TMS pulses delivered at a combination of theta (5Hz) and gamma (50 Hz) frequencies (Huang and Rothwell, 2004). When administered continuously, cTBS has been shown to decrease motor cortical excitability, whereas when delivered intermittently (iTBS), it resulted in motor cortical excitation (Huang et al., 2005). Ozdemir et al. (2021) investigated the reproducibility of the modulatory effects of TBS in the primary motor cortex using concurrent TMS-EEG. Both TBS conditions resulted in substantial variability between and within individuals, with low reproducibility for early and late TEP components (Ozdemir et al., 2021). The reliability of TEPs to assess the neuromodulatory effects of TBS applied to the dorsolateral prefrontal cortex (DLPFC) has not yet been performed. This assessment is important as NIBS is commonly applied to the DLPFC in research and clinical practice as a treatment intervention for many psychiatric and neurological disorders (Schwippel et al., 2019). Showing that the TBS-induced changes in the prefrontal cortical excitability of healthy subjects are reliably using TMS-EEG may help clinical applications to refine stimulation parameters and potentially improve NIBS interventions involving the DLPFC.

The aim of our study is therefore twofold: a) to confirm the reliability of TEP components within an experimental block and between separate sessions as measured by concomitant EEG of single-pulse TMS delivered to the DLPFC; b) to evaluate the reliability of TEPs to measure changes in neural excitability induced by the neuromodulatory effects of iTBS and cTBS applied to the left DLPFC. To this end, the pre-post differences in TEP amplitudes were assessed in different sessions. Comparing the test-retest reliability of TEPs themselves as well as a measure of neural excitability change will help to improve the quality and confidence of future TMS-EEG studies targeting the DLPFC.

## 2. Methods

### 2.1. Participants

Twenty-four right-handed, healthy participants (13 males, mean age 25.2±9.9 years, range from 18 to 65) were included in the study, of which 21 were naïve to TMS. They had to confirm a history of no brain injury or neurological illness, be non-smokers and not use any medication acting on the central nervous system during the study. The Human Research Ethics committee of the University of New South Wales approved the experimental procedures (HC17765). All participants provided written informed consent before the start of the experiment.

### 2.2. Experimental design

Each participant was assessed at five different occasions (2 iTBS sessions, 2 cTBS sessions and one sham session) using a double-blinded crossover design. The order of the visits was pseudo-randomized, with the first three sessions being iTBS, cTBS or sham and the last two sessions iTBS and cTBS, counterbalanced. The EEG responses to 100 single TMS pulses were recorded before each TBS condition (baseline block) and at 2-, 15- and 30-minutes post-TBS (T2, T15 and T30, respectively) (Moffa et al., 2021). All sessions were spaced at least one week apart (to avoid carry-over effects) and scheduled at approximately the same time of the day (to coincide with the same relative time point in each subject’s circadian cycle).

### 2.3. Transcranial magnetic stimulation

Single-pulse stimulation and TBS were administered with a MagPro X100 (MagVenture A/S, Farum, Denmark) stimulator using an MC-B70 figure-of-8 coil oriented 45° to the parasagittal plane using biphasic pulses. The centre of the TMS coil was positioned over the F3 electrode position using a 5-mm customized 3D printed spacer placed between the coil and the scalp to avoid contact with electrodes, minimizing electrical artefacts and electrode movement, as well as bone-conducted auditory and somatosensory inputs. A position was marked on the cap and used as a reference for coil repositioning. A similar procedure was shown to improve consistency in coil location and angle within- and between-sessions when neuronavigation is not available (Rogasch et al., 2013, Chung et al., 2019).

Resting motor threshold (RMT) was determined using the Conforto et al. (2004) procedure by applying TMS over the cap with the spacer. To generate TEPs, participants received 100 single pulses (4s±10% jitter inter-pulse interval) per block delivered at a stimulation intensity of 120% RMT. All TBS interventions were delivered based on the original protocol proposed by Huang et al. (2005): a burst of 3 pulses at 50Hz with 200ms between bursts in a total of 600 pulses. The iTBS protocol involved a 2 s train of TBS repeated every 10 s for a total of 190 s (600 pulses), and cTBS consisted of a 40 s train of uninterrupted TBS (600 pulses).

### 2.4. Electroencephalography (EEG)

EEG data were acquired using a TMSi Refa amplifier (TMS International, Oldenzaal, Netherlands) and 64-channel caps (EASYCAP Gmbh, Herrsching, Germany), as determined by head circumference, with sintered, interrupted disk, Ag-AgCl TMS-compatible electrodes. Electrodes were grounded to Fpz, and EEG signals were measured against an average reference with data sampled at 2048 Hz. To minimise auditory-evoked potentials (AEP) in response to the TMS coil clicking sound, white noise masking was played through in-ear earphones with an extra earmuff placed over them, and volume individually adjusted until the participant could not hear the clicking sound or had reached their upper threshold for comfort.

### 2.5. TMS-EEG processing

All EEG processing and analysis was performed offline using a combination of open-source toolboxes: Fieldtrip (Oostenveld et al., 2011), EEGLAB (Delorme and Makeig, 2004), TESA (v 1.0.1) (Rogasch et al., 2017) and custom scripts on the MATLAB platform (R2017b, The MathWorks, USA). EEG preprocessing followed the same steps as (Moffa et al., 2021), with the cleaning parameters and procedures based on the methods described for the TESA toolbox (Rogasch et al., 2017). Briefly, the main steps were: a) data were epoched between - 1000 to 1000 ms relative to the TMS pulse; b) electrodes with TMS artefact exceeding the input range of the amplifier were excluded and linearly interpolated; c) trials in which kurtosis exceeded 5SD were excluded, and the remaining trials were visually inspected to eliminate those with excessive noise (e.g. muscle activity); d) EEG traces were detrended and corrected relative to pre-TMS data (−500 to −50 ms); e) 50 Hz line noise was removed by fitting and subtracting a sine wave. f) eye blinks were removed with a first round of ICA; e) decay artefacts were removed using a method that fits analytic expressions to the three electrodes with most negative and positive signal deflections to estimate the voltage decay and then subtracts the model fit from the signals (Freche et al., 2018); f) EEG data were filtered using a bandpass filter (1–90Hz); g) With all blocks concatenated, the second round of ICA was performed to remove remaining artefacts (e.g. TMS-muscle, eye movements and electrode noise); h) removed channels were interpolated; i) data were re-referenced to the common average reference.

### 2.6. Data extraction and processing

Our aim was to assess the reliability of the local cortical response to TMS delivered to the left DLPFC. As such, we extracted TEP peak latencies and amplitudes from a region of interest (F1, FC1, F3, and FC3) near the site of stimulation (F3). There is still no consensus on the best way to calculate the peak latencies for extracting the amplitude of TEP components. Therefore, to account for potential differences in latencies between the experimental blocks, amplitude calculations were considered in three different ways: a) Using the grand average TEP for all participants and sessions, including only the pre-TBS blocks to identify component latencies for the assessments of TEP reliability within-block and between-sessions, termed ‘grand base’ latencies henceforth. This was repeated separately now including all blocks, termed hereafter ‘grand all’ latencies for assessments of the reliability of the changes in TEP amplitudes between pre and post-TBS blocks; b) Calculating the TEP per block for each participant by averaging across all sessions, termed ‘block’ latencies; and c) Identifying component latencies for all TEP waveforms, i.e., for each participant per block and per session, termed ‘individual’ latencies. For each of these approaches, peak latencies for TEP components were determined using an algorithm that identified the maximum amplitude for P60 and P200 and the minimum for N40 and N100 within a pre-defined time window (N40: 30–50 ms, P60: 55–75 ms, N100: 90–140 ms and P200: 160–240 ms). When the algorithm failed to identify a particular peak or trough within the defined time window, the amplitude of that component was extracted using the midpoint latency of that component time window (e.g., 40ms for N40, 115ms for N100). The peak amplitude was calculated as the average of values ± 5 ms around the component’s latency. Only experimental blocks with a minimum of 70 trials (after noisy trials were excluded) were included in the reliability analyses. This procedure was adopted to avoid bias when including blocks with a different number of valid trials, as a previous study showed that good concordance could be achieved for most comparisons in the pre-frontal cortex with 70 trials (Kerwin et al., 2018). When a block had more than 70 valid trials, 70 of them were randomly selected to avoid any order effects.

### 2.7. Statistical analysis

A common approach to estimate reliability is the test-retest method which evaluates to what extent the scores arising from a measurement technique are stable for repeated measures over time for the same individual (Mokkink et al., 2010). To this end, data were split (a) within a block, (b) between sessions and (c) as a measure of neuromodulation induced by iTBS and cTBS in separate sessions applying the same active stimulation protocol.

For the within block reliability, comparisons were performed between odd and even trials and between the first and second half of trials obtained from the baseline block of all sessions. The between sessions reliability was assessed by comparing the baseline blocks of all sessions. Finally, the difference in TBS induced neuromodulatory effects between pre/post-stimulation blocks (T2, T15 and T30, respectively 2-, 15- and 30-minutes post-TBS) was calculated considering the two sessions of the same active TBS condition (iTBS or cTBS).

The main statistic used to quantify the reliability of TEPs was the concordance correlation coefficient (CCC), calculated with the following formula (Lawrence and Lin, 1989):

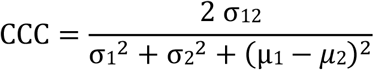

where σ_12_ is the covariance between two partitions, σ_x_^2^ is the variance within a partition, and μ_x_ is the average of the partition.

The choice for CCC over the intra-class correlation coefficient (ICC) was made because the different ICC formulas depend on the assumptions of specific ANOVA models (e.g., normality, homoscedasticity and no multicollinearity) and can be biased if those are not met. In contrast, the CCC is defined without ANOVA assumptions (Chen et al., 2008). The benchmark ranges and labels proposed by Shrout (1998) was used to categorize CCC values into qualitative levels: 0.00-0.10 virtually no reliability, 0.11-0.40 slight, 0.41-0.60 fair, 0.61-0.80 moderate, and 0.81-1.0 substantial reliability.

Finally, to determine whether the CCC values significantly differ from zero, we estimated the 95% confidence interval of the null distribution. TEP amplitudes (for within-block and between-sessions comparisons) or amplitude differences (for the amount of TBS neuromodulation) were randomly permuted for 1000 iterations to determine the CCCs of the permuted data. After checking that the CCC distributions were normal, the 95% CI was determined as ±1.96 times the average standard deviation (SD) of the bootstrapped CCC values of all included trials and latency approaches for that TEP component comparison.

## 3. Results

Data from 24 participants (13 males, mean age 25.2±9.9 years, range from 18 to 65) in a total of 119 sessions were analysed (one cTBS session was excluded due to technical reasons). No participants dropped out during the experiment. The pre-TBS grand-average waveforms revealed peak latencies consistent with previous studies of single-pulse TMS over the DLPFC (Chung et al., 2017, Chung et al., 2019, Conde et al., 2019): N40 (43.5ms), P60 (59.1ms), N100 (117ms) and P200 (202.1ms).

### 3.1 Reliability of TEPs within a block

Regardless of the latency approach used to extract TEP amplitudes, the concordance between odd and even trials of the baseline block showed substantial reliability for N100 and P200 components (CCCs_max_>0.8), whilst P60 showed moderate reliability (0.67<CCC_max_<0.73), and N40 had fair reliability (0.54<CCC_max_<0.56). The CCC values increased with the increasing number of included trials for all components irrespective of the latency approach to extract TEP amplitudes, beginning to plateau after approximately 30 trials (Fig. 1A). CCC values for all components reached their best qualitative reliability category with the fewest trials using the block latencies approach. For example, N100 and P200 achieved substantial concordance after 25 trials with the block latencies but needed almost 35 trials when the grand average latencies were used (supplementary Table 1). N40 and P60 reached their maximum concordance with even fewer trials (approximately 15), although achieving inferior qualitative levels in comparison to the later components even when all 35 trials were compared (fair and moderate, respectively).

**Fig 1.**
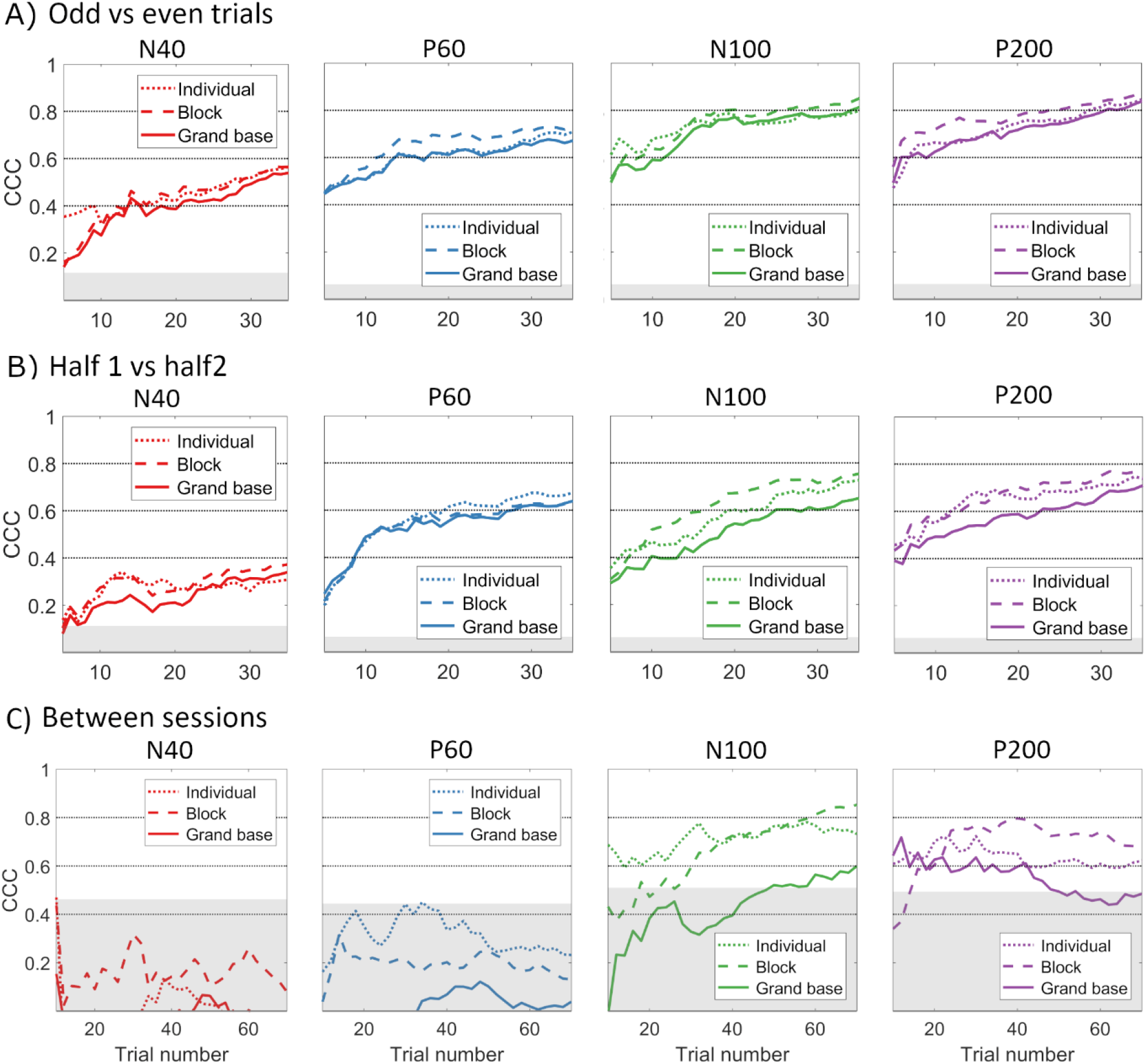
Concordance of TMS-evoked potentials within and between the baseline blocks. A) Concordance correlation coefficient between odds and even trials of baseline block. B) Concordance correlation coefficient between the first and second half of the baseline block. C) Concordance correlation coefficient between the baseline block across five separate sessions. Shaded regions in grey show the 95% CI of the null distribution. The three lines show the different latency strategies used for amplitude extraction, and the x-axis shows the number of trials used for estimating the CCC. Horizontal lines at 0.4, 0.6 and 0.8 are the lower limits for the fair, moderate and substantial reliability categories, respectively.

When comparing TEPs calculated with the trials of the first half versus the second half of the baseline block, P60, N100, and P200 presented similar CCC qualitative level (moderate reliability) for all latency approaches, with numerical results of P200 (0.71<CCC_max_<0.78) slightly superior to the CCC of N100 (0.65<CCC_max_<0.75) and P60 (0.64<CCC_max_<0.67). The N40 component again presented the lowest CCCs achieving a slight reliability level (0.34<CCC_max_<0.37). Once again, the CCC values showed an increasing trend with the growing number of trials for all components, with N40 and P60 plateauing after 30 trials, whereas N100 and P200 maintained growth (Fig. 1B). TEP amplitudes extracted with the block latencies provided the higher numerical CCCs for the later components, achieving a higher reliability category with fewer trials (around 15 for both N100 and P200) (supplementary Table 2). In contrast, for the P60, the individualised method led the CCC values to the moderate category with fewer trials (20) than the other approaches. Finally, regardless of the latency method, N40 did not exceed the range of fair concordance even with all 35 trials considered.

### 3.2 Reliability of TEPs between sessions

The reliability estimations for the pre-TBS block across all five experimental sessions showed that the N100 component reached the highest concordance (CCC_max_=0.85), achieving substantial reliability with the block-latencies amplitudes (Fig. 1C). A moderate concordance was achieved by P200 (CCC_max_=0.68), also using the same amplitude extraction method. For both components, the grand-latencies approach led to the lowest CCCs providing a fair reliability level (CCC_max_=0.6 for N100 and CCC_max_=0.48 for P200), while the extraction of amplitudes using the individualised latency resulted in a moderate reliability level (CCC_max_=0.73 for N100 and CCC_max_=0.62 for P200).

The CCC increasing trend with the number of trials was observed for the between-sessions comparisons of the N100 component, which needed 60 trials to achieve substantial concordance when the block latencies were used (Fig. 1C). This increasing trend was also observed for the P200 until around 30 trials when the individualised and block latencies strategies were used, after which CCC values stabilized within the moderate concordance range. For the grand average approach, P200 CCCs fluctuated up to 40 trials between moderate and fair concordance levels, after which the reliability level showed a downward trend and was not significantly different from the null distribution when all 70 trials were analysed.

CCC reliability for N40 and P60 did not significantly exceed the 95% CI of the null distribution for any of the amplitude extraction methods, even considering the maximum number of trials.

### 3.3 Reliability of the neuromodulatory effects induced by iTBS and cTBS

Regardless of the latency methodology used to extract TEP amplitudes, the block T2 starting 2 minutes after the end of iTBS presented the lowest concordance levels for the change in amplitudes for all components (Fig. 2A and supplementary figure 2). In general, the N100 was the component showing the highest concordance between the changes in amplitude after iTBS across repeated sessions.

**Figure 2.**
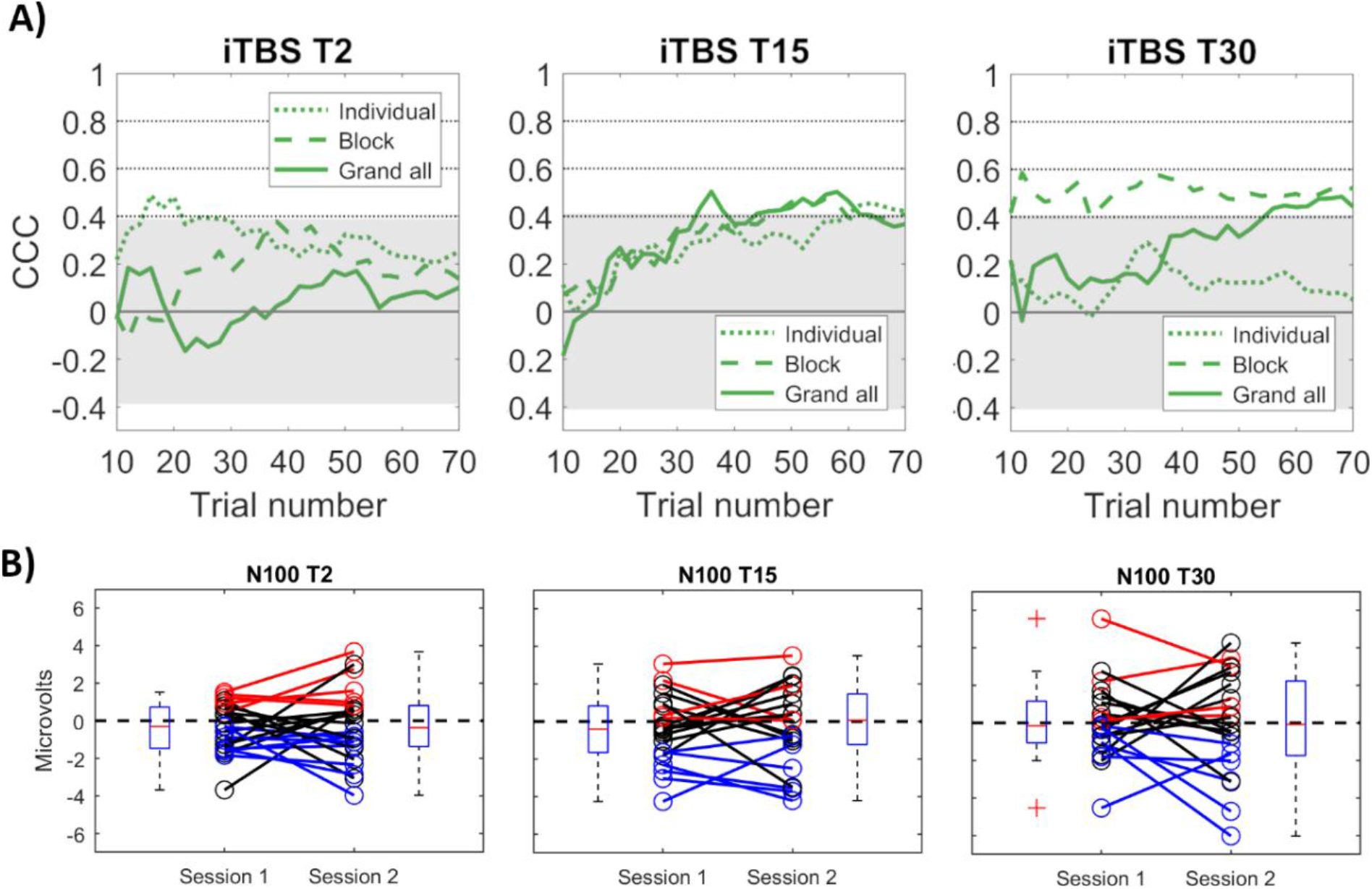
Neuromodulatory effects induced by iTBS on the N100 component. A) Concordance correlation coefficients (CCC) of the amount of neuromodulatory effects induced by iTBS between each post-iTBS block minus pre-across trial numbers and latency strategies used for amplitude extraction. B) Difference in amplitudes of TMS-evoked potentials between post-iTBS and pre and boxplot for visits 1 and 2. Lines in red represent pairs of observations for each participant in which the difference in amplitudes had positive values in both visits. Lines in blue represent pairs of observations in which the difference in amplitudes had negative values in both visits. Lines in black represent pairs of observations in which the difference in amplitudes had opposite signs between the two visits.

The concordance of the change in N100 amplitude increased with the number of trials for all latency approaches, reaching fair reliability at T15 with block and individual latencies (CCC_max_ = 0.42) and T30 with the block latencies (CCC_max_=0.52) and amplitudes extracted using grand all-latencies (CCC_max_ =0.44) (Fig. 2A and supplementary Table 4).

The change in P200 achieved fair reliability (CCC_max_=0.47) only at T15 with approximately 30 trials, using the amplitudes extracted by the grand all latencies method (supplementary Fig. S3). For the changes in P60, CCC measures at 30 minutes post-iTBS showed an increasing trend up to approximately 35 trials, varying after that until stabilizing to achieve with 70 trials a marginally significant slight concordance (CCC_max_=0.40) with the use of the individual and block latency amplitudes (supplementary Fig. S3). Finally, N40 did not exceed the 95% CI of the CCCs’ null distribution in any experimental blocks’ post-iTBS regardless of the latency approach used to extract TEP amplitudes.

In contrast, the changes in TEP amplitudes after cTBS achieved a significant fair reliability level only for P60 at T2 when the individualized latencies methodology was used (CCC_max_=0.50), with all other components’ CCCs below the 95% CI of the null distribution (supplementary Fig. S4).

The magnitude and direction of iTBS and cTBS induced changes on TEP amplitudes showed large inter-individual variability (supplementary Fig. S6-S11). For example, the change in N100 amplitudes after iTBS for the first and second experimental sessions using block latencies is summarized in Fig. 2B. The proportion of participants that presented a more negative N100 amplitude after iTBS in both visits was 58%, 46% and 52%, respectively at T2, T15 and T30. A smaller number of participants presented opposite directions (increase/decrease) on the change of N100 across the two visits: 4%, 18% and 5%, at T2, T15 and T30, respectively.

### 3.4 Comparison of TEP’s reliability measures

Finally, we compared the test-retest reliability of TEPS within and between sessions and for detecting the neuromodulatory effects of TBS. In general, the number of trials that each component needed for the same qualitative reliability level increased with the duration between repeated assessments, i.e., from within block to between sessions. Likewise, the reliability achieved with the same number of trials (35) decreased with the duration between repeated assessments (Fig. 3A). This decreasing trend was more pronounced for the early components N40 and P60, which showed a greater reduction in reliability when assessed in different sessions, only achieving slight reliability between sessions. In contrast, the reliability of the N100 and P200 components remained fairly stable when assessed in different sessions using the block latencies methodology (Fig. 3A). We observed the lowest CCC levels for the comparisons between separate sessions when using the grand average baseline latencies, impacting all components, particularly N40 and P60, which showed a sharp reduction in concordance compared to within-block assessments (supplementary Fig. S1B).

**Figure 3.**
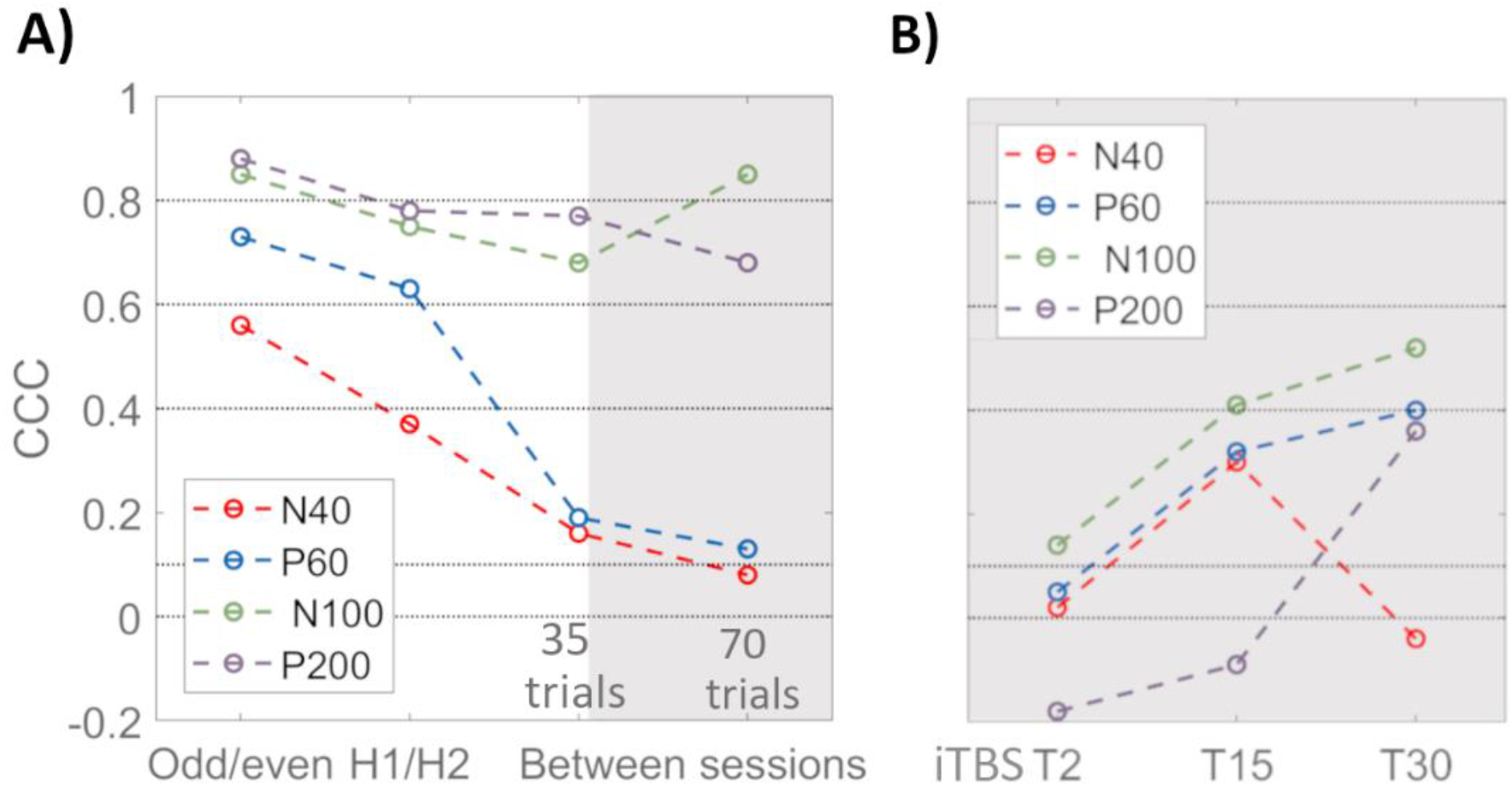
The concordance correlation coefficient for each component using TEP amplitudes extracted based on the block latencies. A) CCC is plotted for the three levels of comparisons: within the baseline block (odd vs even and half 1 vs half 2) and between the baseline block across five separate sessions. All CCC values shown in panel A are calculated for 35 trials. B) CCCs for the change in TEP amplitudes from 2 to 15 and 30 minutes post-iTBS (T2, T15 and T30, respectively). Horizontal lines at 0.1, 0.4, 0.6 and 0.8 are the lower limits for the slight, fair, moderate, and substantial reliability categories, respectively. Each TEP component is colour coded according to the legend inside the figures.

The concordance was generally lower when assessing the neuromodulatory effects of TBS. The highest concordance was observed for the N100 following iTBS, which showed an increase with time to fair concordance at 30 minutes after iTBS (Fig. 3B). Increasing concordance with time after iTBS was also observed for the P60 and P200 components. However, for the P60 component, the reliability increased above the level of concordance observed for the TEPs assessed in different sessions (Fig 3A), questioning whether this increase was genuine.

When comparing the different methods to determine the latencies of the TEP components, using the block latencies to extract TEP amplitudes resulted in higher concordance values in most reliability assessments for all components. The only exception was P60 which showed higher concordance with the individual latencies approach for the comparison between sessions (CCC=0.42 vs CCC=0.19 for the block latencies) (supplementary Fig. 1A and Fig. 3A, respectively). The influence of the latency method choice in reliability outcomes was more pronounced for comparisons between sessions than for within-block analyses (Fig. 1).

## 4. Discussion

We investigated the test-retest reliability of TMS-evoked potentials measured at the DLPFC both before and after repeated sessions of iTBS and cTBS in healthy participants. Our results showed that overall, the TEP peaks with the best within-block reliability were P60, N100 and P200, while only N100 and P200 presented good concordance between sessions at baseline. The N40 presented relative lower levels of concordance within-block and was not reliable for baseline between-sessions comparisons. TBS-induced effects on cortical excitability only reached moderate reliability (CCC_max_ of 0.52) for the N100 amplitude assessed at T30 after iTBS. The concordance for all other components and time points was slight to fair. Hence, the present findings show that while the reliability of the TEPs themselves was fairly good, particularly for the later N100 and P200 components, the reliability of TEPs to detect changes in neural excitability induced by iTBS and cTBS was limited. That is, in general, measures are only considered adequate when the substantial reliability category is achieved (Shrout, 1998).

### Reliability of TEPs within-block and between sessions

Both N100 and P200 components achieved substantial reliability when comparing odd and even trials, and moderate reliability for comparisons between the two halves of the pre-TBS block, regardless of the latency strategy used to extract the component amplitudes. When comparing the pre-stimulation (baseline) block of the five different sessions, the concordance of the N100 was marginally better, achieving substantial reliability, while the P200 showed moderate agreement. In general, reliability estimates within the same session were higher than between different sessions for the N40, P60 and P200 components, whereas the N100 maintained similar levels across the different data split comparisons. Moreover, the N40 and P60 components were consistently less reliable than the later components (N100 and P200) for comparisons within the baseline block and did not achieve significant reliability between sessions.

The observed high test-retest concordance levels of N100 and P200 within and between sessions are consistent with a previous study on the DLPFC showing similar reliability levels for the prefrontal ROI (DLPFC) when the same number of trials were considered (Kerwin et al., 2018), suggesting that these components are the most robust candidates to be explored in future experimental and clinical studies. However, it is believed that the time window encompassing the N100 and P200 components are also the most susceptible to contamination by somatosensory- and auditory-evoked potentials (Rocchi et al., 2021), which can still be present despite applying state-of-the-art procedures to minimise this type of peripheral input. Therefore, peripheral-evoked potentials may contribute to a relatively more stereotypical response to TMS in the time window encompassing the N100 and P200 (Conde et al., 2019), leading to higher CCC levels. Nonetheless, TMS-EEG experiments performed on the motor cortex using a ‘realistic’ sham procedure as a control condition suggest that N100 does carry a genuine neural signal of TMS effects (Gordon et al., 2021, Gordon et al., 2018). Another TMS-EEG study of the motor cortex with a design to separately estimate the contribution of AEP and SEP induced by TMS show that, if the procedures to minimise indirect cortical activation are properly adopted, it is possible to disentangle genuine cortical activity from the EEG signals (Rocchi et al., 2021).

On the other hand, the N40 and P60 peaks were unreliable for comparisons between separate days, which deviates from the first study assessing the reliability of repeated measures with correlation coefficients (Lioumis et al., 2009). However, this finding is in line with a more recent investigation that showed lower concordance of these early components amplitudes on separate days (CCC_max_<0.6) in a region of interest encompassing the left prefrontal cortex (Kerwin et al., 2018). This discrepancy may be explained because correlation coefficients are not considered the ideal metric to assess test-retest reliability (Koo and Li, 2016). The time window encompassing N40 (30-50 ms) can be affected by voltage decay artefacts induced by the magnetic field interaction with the electrode-skin interface (Veniero et al., 2009) and cables (Sekiguchi et al., 2011), or caused by the tail of muscle activity (Rogasch et al., 2014), in particular, due to the activation of cranial and facial muscles from a more lateralised stimulation site like F3 (Huang and Mouraux, 2015, Ilmoniemi and Kičić, 2010). Even though a mix of ICA (Rogasch et al., 2017) and subtraction of fitting analytical expressions (Freche et al., 2018) was used during the offline postprocessing, the strongly induced muscle twitches typical of prefrontal stimulation with TMS may have superseded the capacity of the cleaning procedures to fully remove these artifacts without attenuating the genuine neural signals (Rogasch et al., 2021). This may have contributed to the relatively lower CCC values of earlier components, imposing challenges and limitations on using N40 as potential biomarker in multi-session studies stimulating a highly artefactual region such as the DLPFC.

### Reliability of the change in TEP amplitudes after iTBS and cTBS

Though early TEP components (15-55 ms) are suggested to reflect the cortical activity close to the stimulation site (Rogasch et al., 2020), a marginal (i.e., ‘fair’) reliability level of the change in amplitudes from pre to post iTBS was observed only for the N100, P200 and P60 components at specific time points.

Even though the within-block and between-session comparisons for N100 were substantially reliable and reproducible, iTBS induced changes on the N100 amplitude presented only fair reproducibility across sessions and were highly variable intra- and inter subjects at all experimental time points (Fig. 3B and supplementary Fig.S6-S11). Fair reliability means that approximately half of the observed variance can be attributed to measurement error (Matheson, 2019). The limited concordance of TBS-induced changes between sessions may indicate that TEPs are not a reliable marker of changes in cortical excitability or reflect a lack of consistent effects on prefrontal cortical excitability induced by TBS. Similar variability has also been observed in the motor cortex when assessed with TEPs (Ozdemir et al., 2021) and MEPs (Corp et al., 2020). The reasons underlying such variability in response to the stimulation remains unclear. However, a possible explanation may lie in a combination of trait related and state-dependent causes (Ziemann and Siebner, 2015). For example, differences in the polymorphism of the brain-derived neurotrophic factor (BDNF) gene were shown to impact the plasticity induced by iTBS in the motor cortex (Antal et al., 2010). The underlying physiological brain activity at the stimulation time has also been considered to critically influence the changes in cortical excitability induced by rTMS protocols (Romei et al., 2007, Huang et al., 2017).

For cTBS, P60 was the only component that exceeded the 95% CI of the null distribution, reaching fair concordance in the first block post-stimulation for amplitudes extracted with the individualised latency. Meta-analytical data showed that cTBS applied to the motor cortex led to a decrease in corticospinal excitability with the greatest effect sizes at early time points post-stimulation (≤ 5 min) (Chung et al., 2016). However, in our previous study using a subset of the present sessions (the first iTBS/cTBS and sham sessions), no significant differences in the amplitudes of TEPs were found between cTBS, iTBS and sham stimulation (Moffa et al., 2021). In addition, the reliability was above the level of concordance observed for the P60 assessed in different sessions (Fig 3A), hence questioning whether the observed fair concordance of the P60 reflects a genuine effect. That is, the limited reliability of N40 and P60 on the DLPFC puts an upper boundary on the utility of these components and may have diminished the power of the study to find true differences in local neuromodulatory effects between conditions that they could potentially reflect (Moffa et al., 2021).

Several variables may contribute to between-sessions variability, which is likely to impact the test-retest reliability levels of TBS effects. Past evidence suggests that TEPs are sensitive to variations in brain state (Peters et al., 2020), level of fatigue (Otieno et al., 2019), vigilance state (Massimini et al., 2005) and movement initiation (Nikulin et al., 2003). The level of vigilance with time awake from the morning also seems to impact cortical responsiveness to TMS (Manganotti et al., 2013, Huber et al., 2013). We, therefore, took various precautions in the design and conduction of TMS-EEG experiments to minimise the influence of these factors. For example, the sessions were conducted approximately at the same time of day to account for each individual’s circadian cycle, and we instructed the participant to stay relaxed but not sleep during the procedures. Variations in vigilance state and fatigue may also impact the latencies of the TEP components (Massimini et al., 2005, Otieno et al., 2019), which may further impact the reliability of TEPs between blocks depending on the method used for defining peaks latencies. Using the participant block latency to extract component amplitudes in our dataset resulted in higher concordance levels for most reliability assessments. The choice of the participant block method showed greater influence when comparing sessions from different days, overcoming the relatively lower specificity provided by the grand average and the likely greater variability and lower signal-to-noise ratio of the individualised latency approach, which uses a smaller number of trials in its estimation. In contrast, all latency strategies led to similar CCC levels for the within-block evaluations, suggesting this aspect is not as crucial as in assessments involving multiple sessions.

## 5. Limitations

We did not use neuronavigation to position the TMS coil. Although marks in the EEG caps can consistently guide TMS coil repositioning over the target region (Rogasch et al., 2013, Chung et al., 2019), it does not control for coil tangentiality, which might have increased measurement error between blocks and sessions due to potential variations in the strength of the induced electric field delivered to the target region (Casarotto et al., 2010). We quantified TEP peaks at only one region of interest encompassing the left DLPFC, as we were interested in the reliability of the local effects of TBS. A previous study also stimulating the DLPFC observed the highest reliability for some comparisons in ROIs related to other brain regions, e.g., the central and parietal electrodes for later evoked potentials (Kerwin et al., 2018). Therefore, the reliability values will probably differ when estimated in other brain regions. Due to its more anterior and lateral position and the proximity to facial muscles and cranial nerves, the stimulation of the DLPFC tends to result in a large artefactual signal that could have impacted the reliability of the earlier components like the N40 (Rogasch et al., 2013). The reliability estimations were performed, including blocks with more than 30 bad were excluded. The CCC values generally showed an increasing trend with increasing number of trials. Therefore, it could be possible that increasing the number of accepted trials could lead to higher TEP reliability levels. The present results are related to healthy volunteers with the specific demographics of the sample. Therefore, extrapolating the present results to clinical populations should be done with caution. Finally, the amplitude and reliability of TEPs from the DLPFC are dependent on the processing pipeline (Bertazzoli et al., 2021), and a consensus on the processing pipeline has not yet been established.

## 6. Conclusion

This study reports the first test-retest examination of the reliability of the neuromodulatory effects of TBS over the DLPFC using TMS-EEG. TBS-induced changes on TEP amplitudes indicated only fair reproducibility levels for P60, N100 and P200 at specific time points, and no reliable changes after cTBS. Only the N100 and P200 achieved a good level of concordance within a block and across separate sessions and are the best options to be considered in future experimental and clinical studies. Lower concordance of the earlier components could reflect residual TMS-induced artefacts, limiting their utility in assessing the DLPFC with multiple visits. The neuromodulatory effects of TBS protocols on cortical excitability were marked with substantial between- and within-subjects variability, indicating the importance of designing experiments with repeated assessments to validate neuromodulatory findings and the use of increased sample sizes to overcome this variance. These findings suggest that individual or group results of TBS modulation effects based on a single assessment need to be considered with caution.

Additional studies with greater sample size and realistic control condition methods are needed to further clarify the sources of variability in response to TBS and improve the reliability of TEPs. Finally, sharing data, codes, and procedures among the TMS-EEG community is paramount to validate findings, techniques and methods, and to aim for a consensus of the best online and offline procedures to measure TEPs that reliably estimate the neuromodulatory effects of NIBS techniques and that could aid in the refinement of these interventions for the treatment of neuropsychiatric disorders. With this in mind, the data reported in this manuscript will be made publically available for interested researchers.

## Acknowledgements

The authors wish to thank Nichola Jephcott, Design Futures Lab, School of Built Environment, UNSW, for the assistance with the customised 3D printed spacer, and all volunteers for their participation. A.H. Moffa was the recipient of a Scientia PhD Scholarship from the University of New South Wales, Sydney, Australia and received support through an “Australian Government Research Training Program Scholarship”. The present work will contribute to the doctoral thesis of A. H. Moffa.

