## Supplementary Material for "Reliability of transcranial magnetic stimulation evoked potentials to detect the effects of theta-burst stimulation of the prefrontal cortex"

**Sup. Table1.** Reliability of TEP components (CCC) within the baseline block with comparisons performed between odd and even trials. Each column is related to the specif latency approach used to extract TEP amplitude. Colours refer to the benchmark values and labels proposed by Shrout (1998).

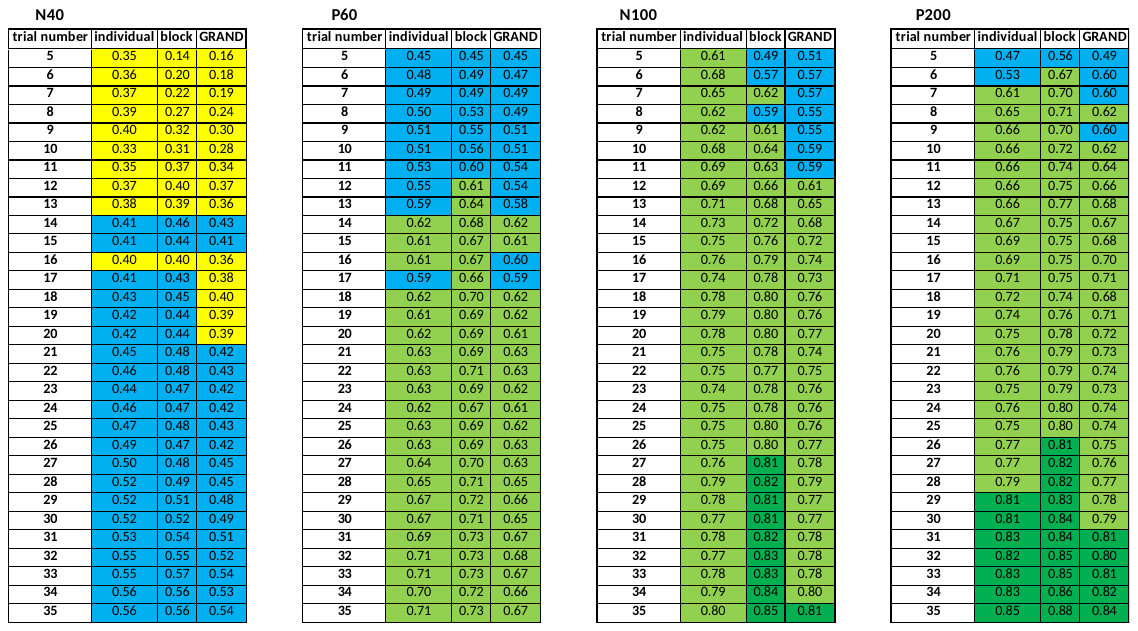

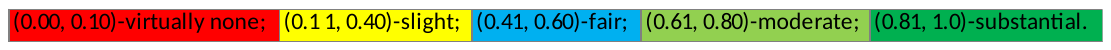

**Sup. Table2.** Reliability of TEP components (CCC) within the baseline block with comparisons performed between the first and second half of the trials. Each column is related to the specif latency approach used to extract TEP amplitude. Colours refer to the benchmark values and labels proposed by Shrout (1998)

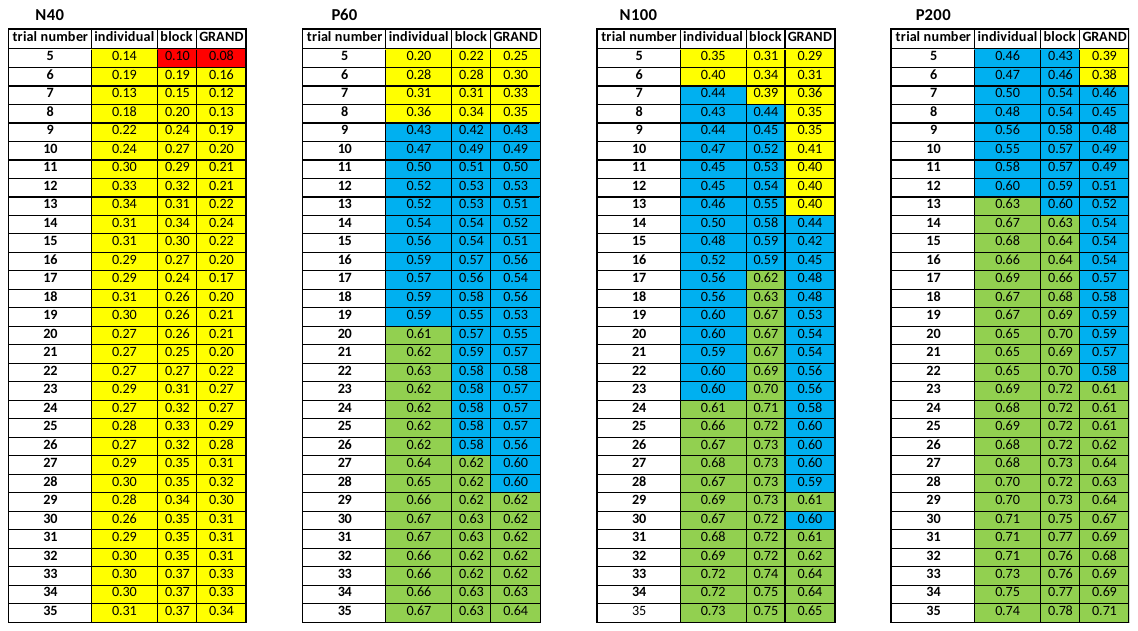

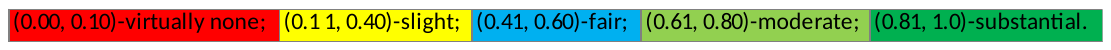

**Sup. Table3.** Reliability of TEP components (CCC) between the baseline block of five separate sessions. Each column is related to the specif latency approach used to extract TEP amplitude. Colours refer to the benchmark values and labels proposed by Shrout (1998).

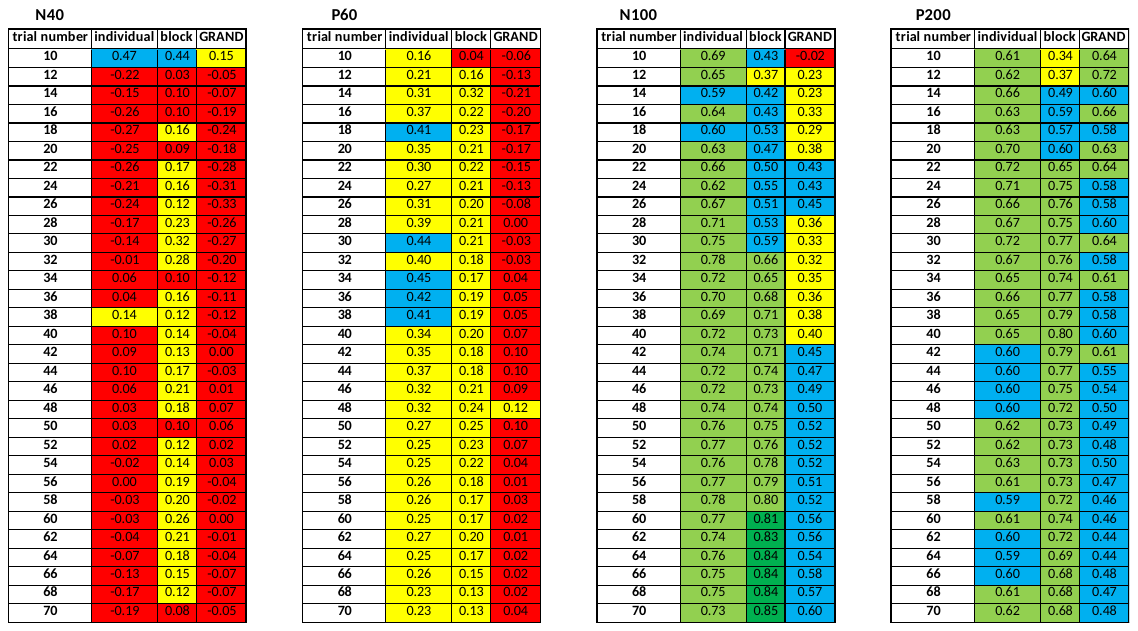

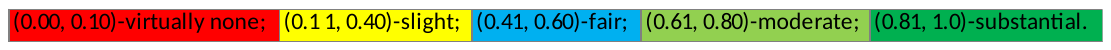

**Figure S1**. The concordance correlation coefficient for each TEP component is plotted for the three levels of comparisons for each component (colour coded legend inside figures). CCCs for all comparisons are plotted for 35 trials. A) Using ‘individual latencies’; B) Using ‘grand average pre-latencies’.

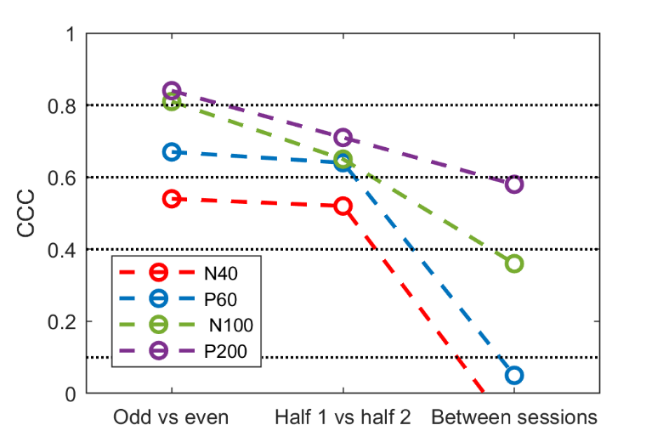
A) **Individual latencies**	B) **Grand base latencies**

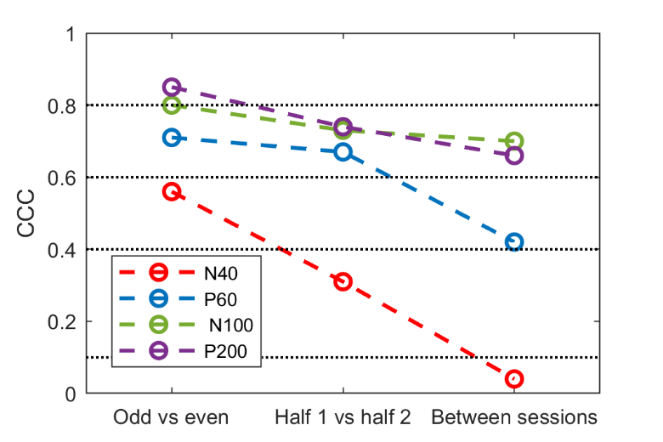

**Figure S2**. The concordance correlation coefficient for the changes in TEP amplitudes 2, 15 and 30 minutes after iTBS (T2, T15 and T30, respectively) for each component (colour coded legend inside figures) for the three latency methodologies used to extract TEP amplitudes. Colours refer to the benchmark values and labels proposed by Shrout (1998). CCCs for all comparisons are plotted for 70 trials. A) Using ‘individual latencies’; B) Using ‘grand average pre-latencies’.

A) **Individual latencies**	B) **Grand all latencies**

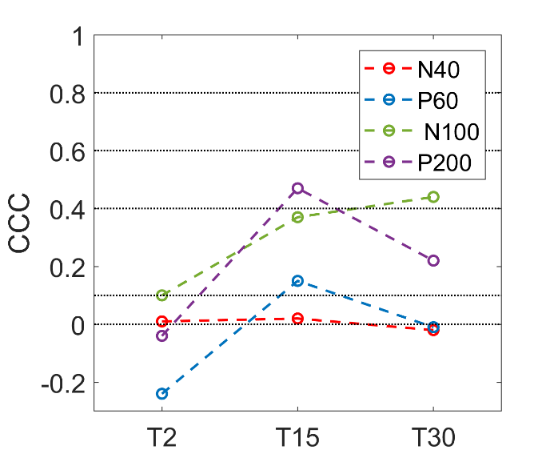

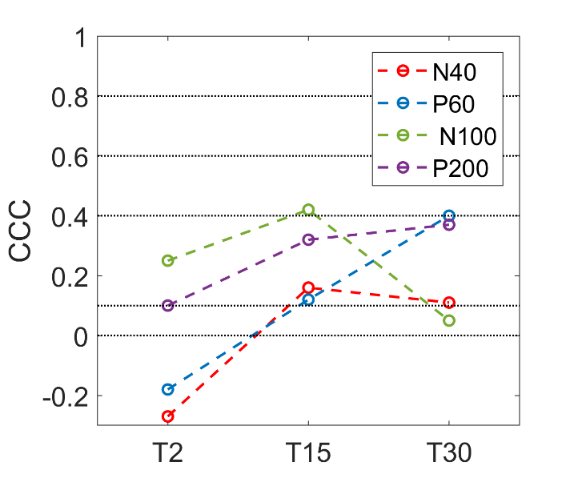

**Sup. Table 4.** Concordance correlation coefficient (CCC) for the changes in TEP amplitudes 2, 15 and 30 minutes after iTBS and cTBS (T2, T15 and T30, respectively) for the three latency methodologies used to extract TEP amplitudes. Colours refer to the benchmark values and labels proposed by Shrout (1998).

(0.00, 0.10)-virtually none;

(0.1 1, 0.40)-slight;

(0.41, 0.60)-fair;

(0.61, 0.80)-moderate;

(0.81, 1.0)-substantial.

**A) Grand all latencies**

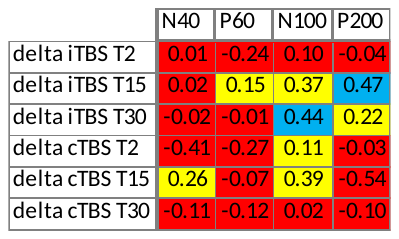

**B) Block latencies**

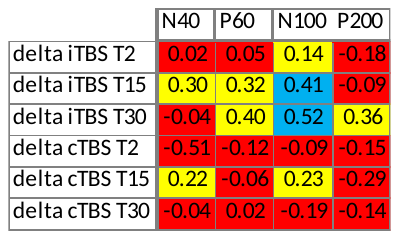

**C) Individual latencies**

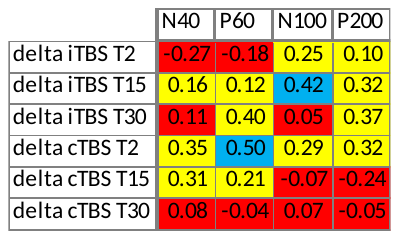

**Figure S3.** N**euromodulatory effects induced by iTBS.** Concordance correlation coefficients (CCC) of the amount of neuromodulatory effects induced by iTBS between each post-iTBS block minus pre-iTBS across trial numbers and latency strategies used for amplitude extraction.

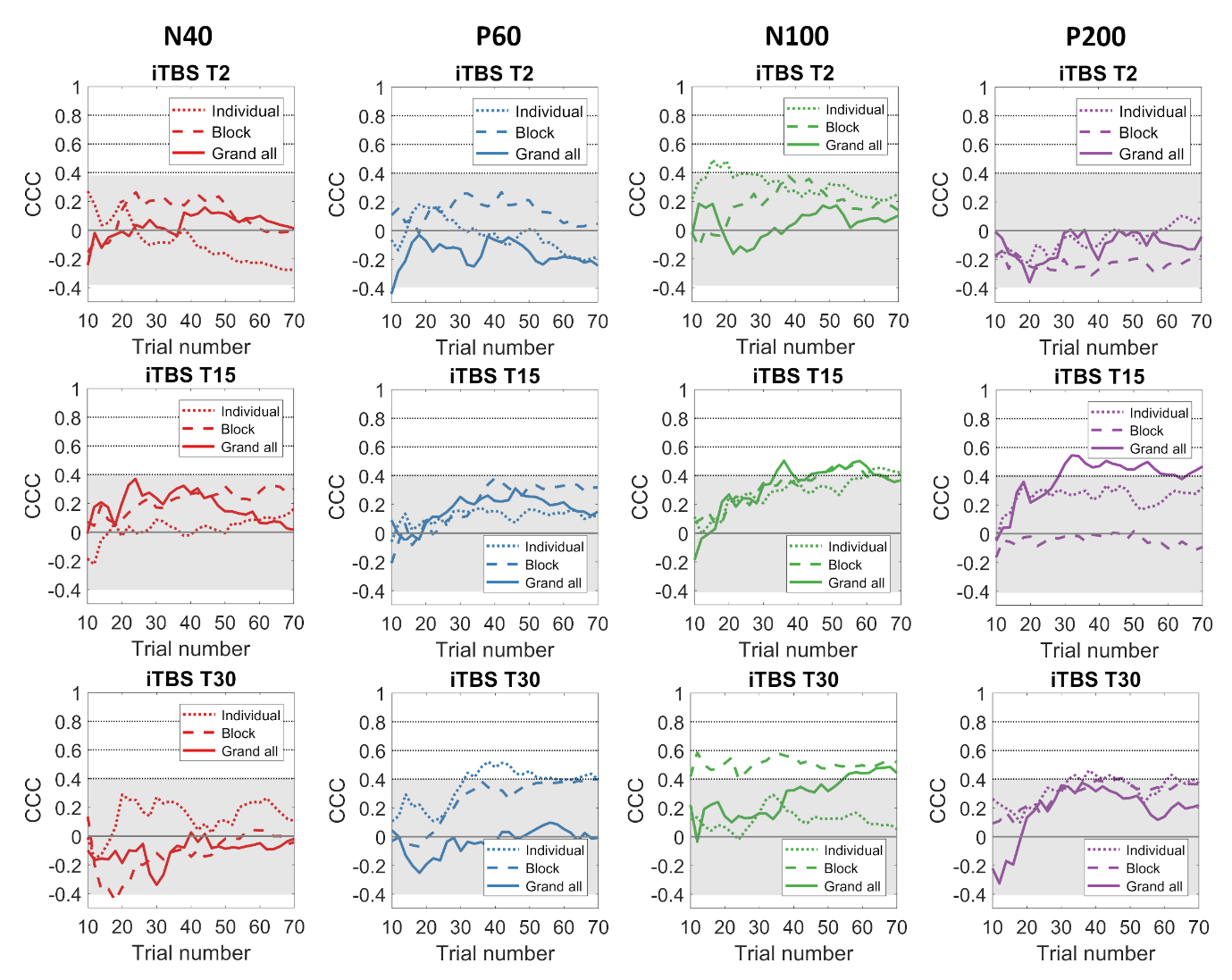

**Figure S5. Neuromodulatory effects induced by cTBS** Concordance correlation coefficients (CCC) of the amount of neuromodulatory effects induced by cTBS between each of the post-cTBS blocks minus pre-cTBS across trial numbers and latency strategies used for amplitude extraction.

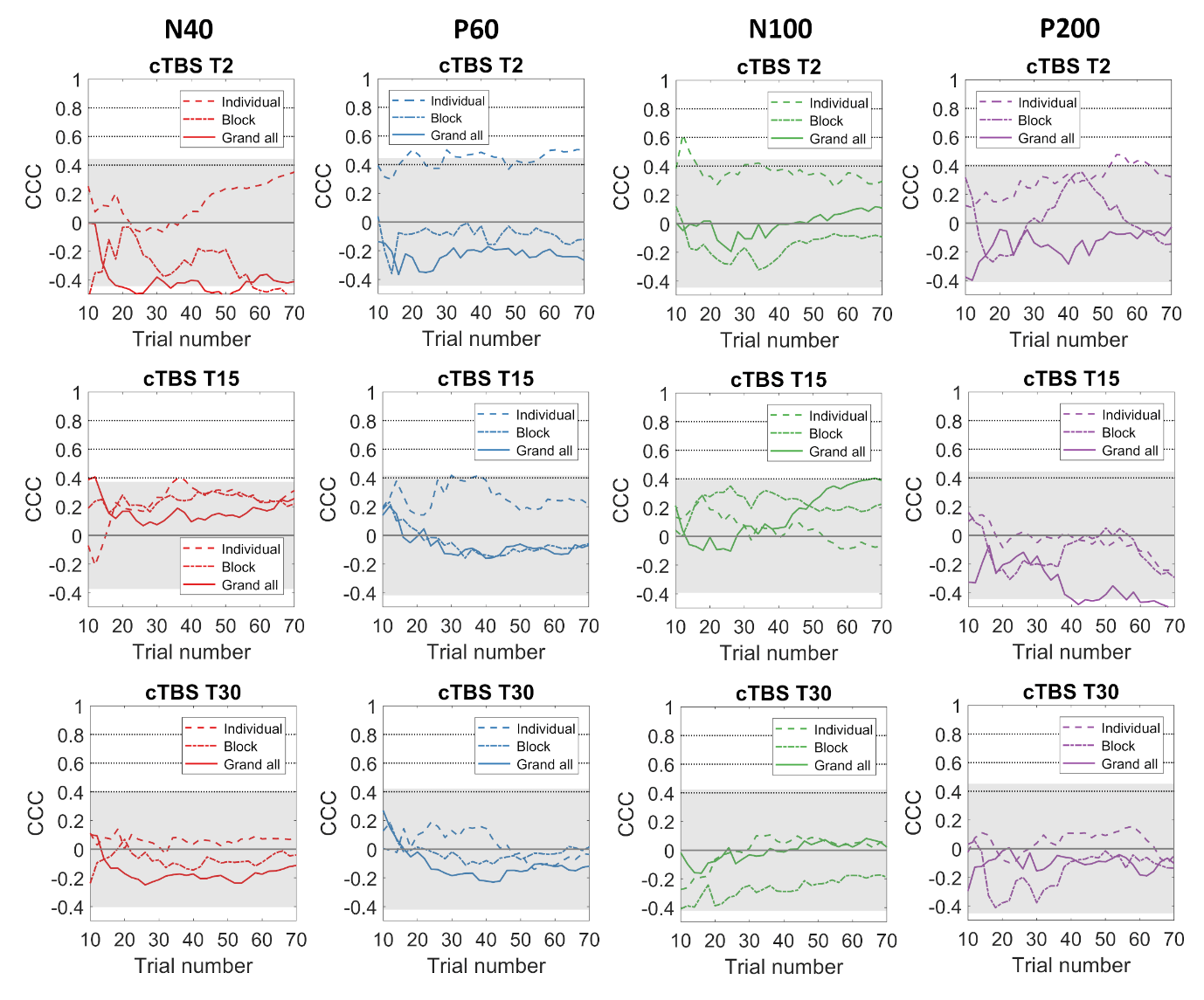

**Figure S6-S8.** Differences in amplitudes of TMS-evoked potentials between post-iTBS (T2, T15 and T30) and pre and boxplot for visits 1 and 2 with grand average, participant block and individualised latency approaches. Lines in red represent pairs of observations for each participant in which the difference in amplitudes had positive values in both visits. Lines in blue represent pairs of observations in which the difference in amplitudes had negative values in both visits. Lines in black represent pairs of observations in which the difference in amplitudes had opposite signs between the two visits.

**Figure S6. Using ‘grand all’ latencies**
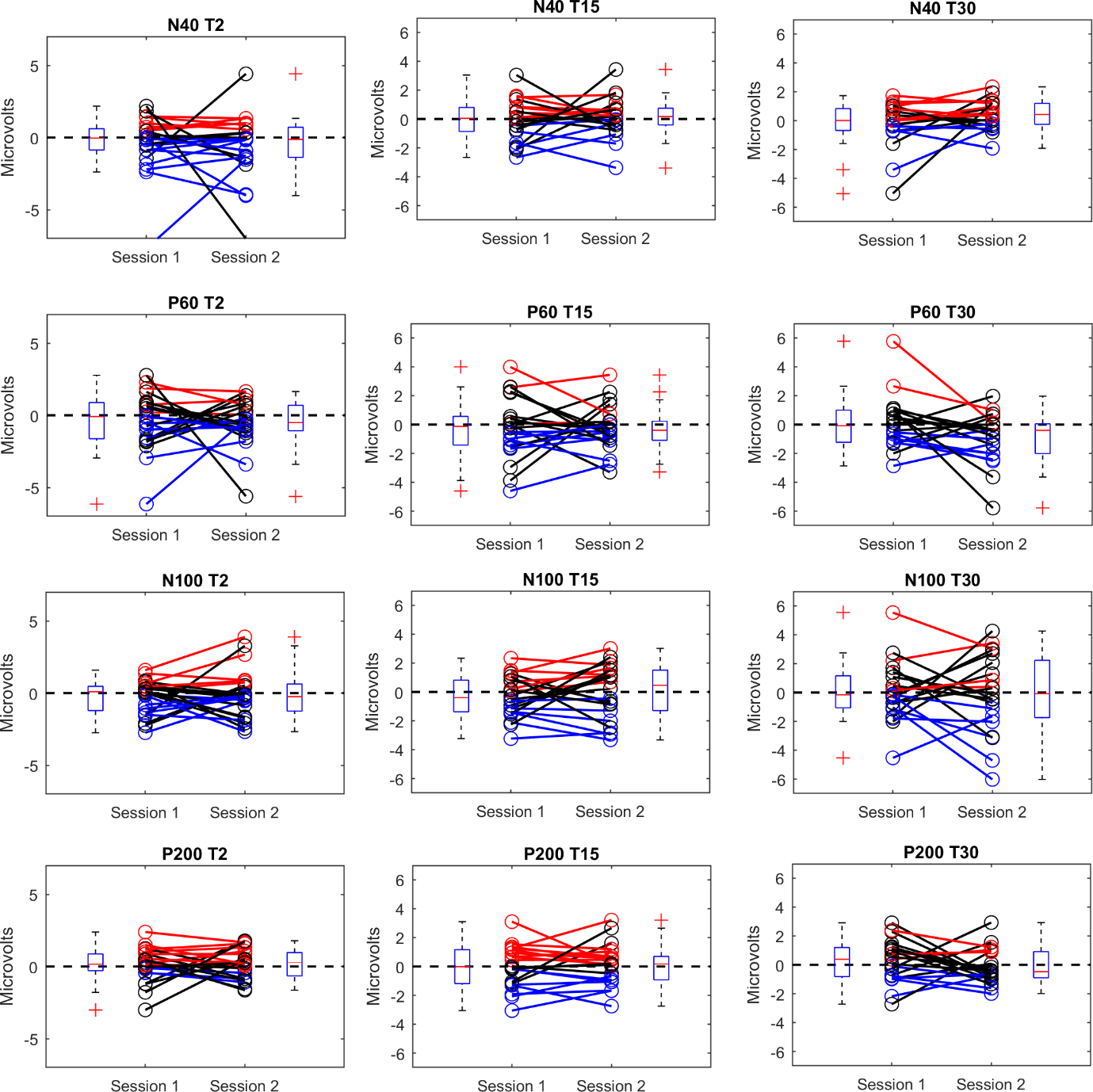

**Figure S7. Using ‘block’ latencies**

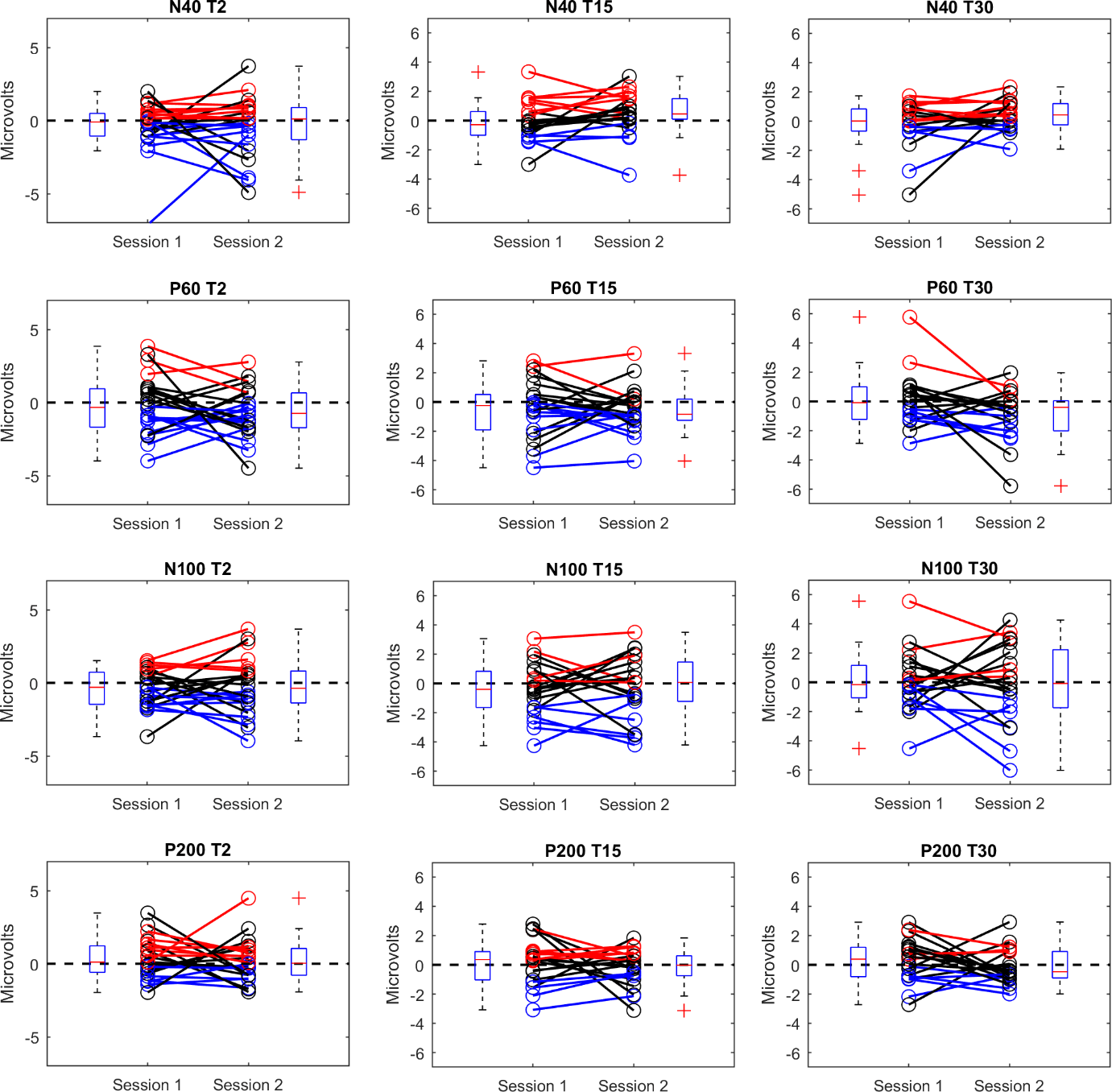

**Figure S8. Using ‘individual’ latencies**

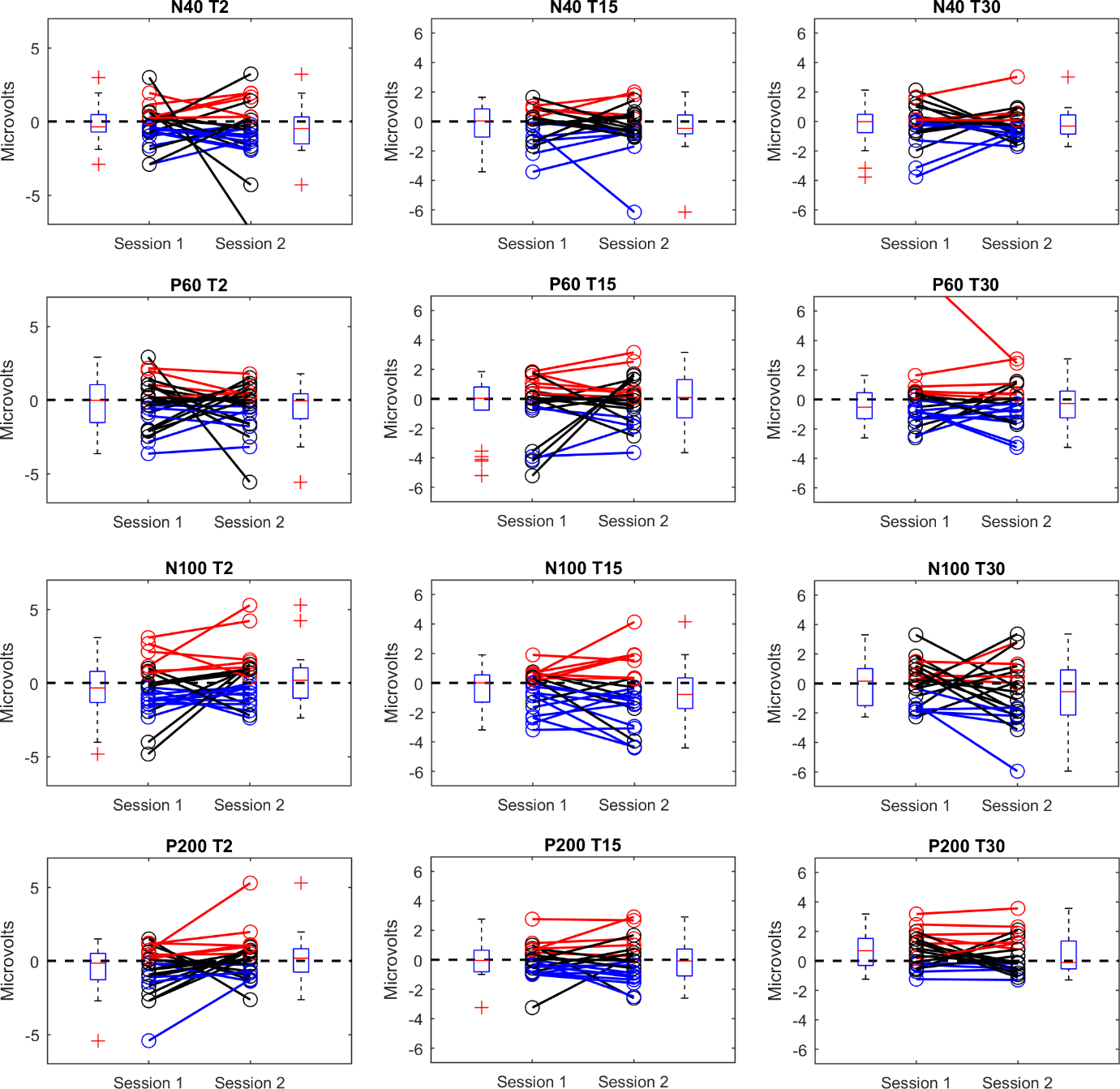

**Figure S9-11.** Differences in amplitudes of TMS-evoked potentials between post-cTBS (T2, T15 and T30) and pre and boxplot for visits 1 and 2 with grand average, participant block and individualised latency approaches. Lines in red represent pairs of observations for each participant in which the difference in amplitudes had positive values in both visits. Lines in blue represent pairs of observations in which the difference in amplitudes had negative values in both visits. Lines in black represent pairs of observations in which the difference in amplitudes had opposite signs between the two visits.

**Figure S9. Using ‘grand all’ latencies**

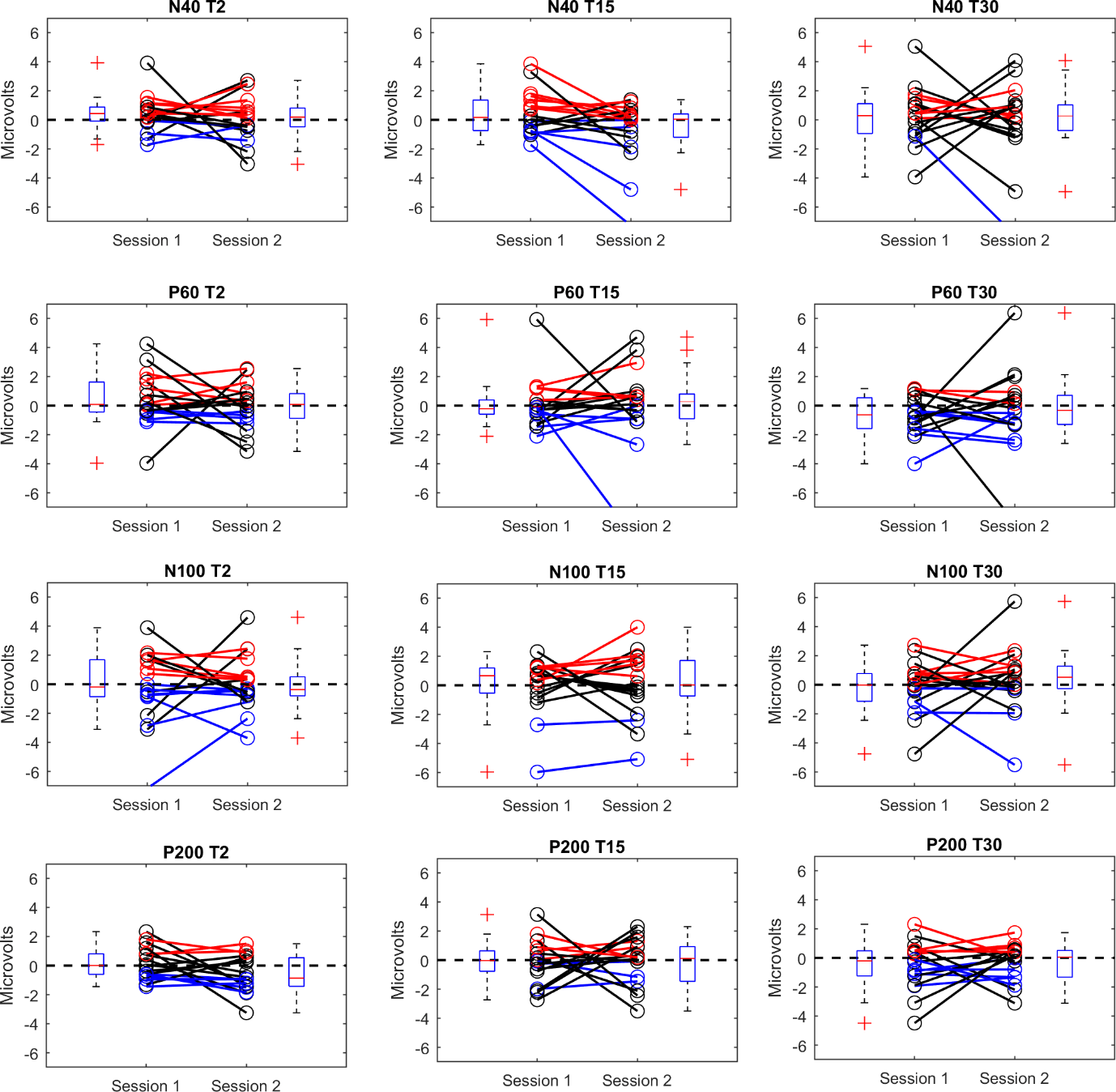

**Figure S10. Using ‘block’ latencies**

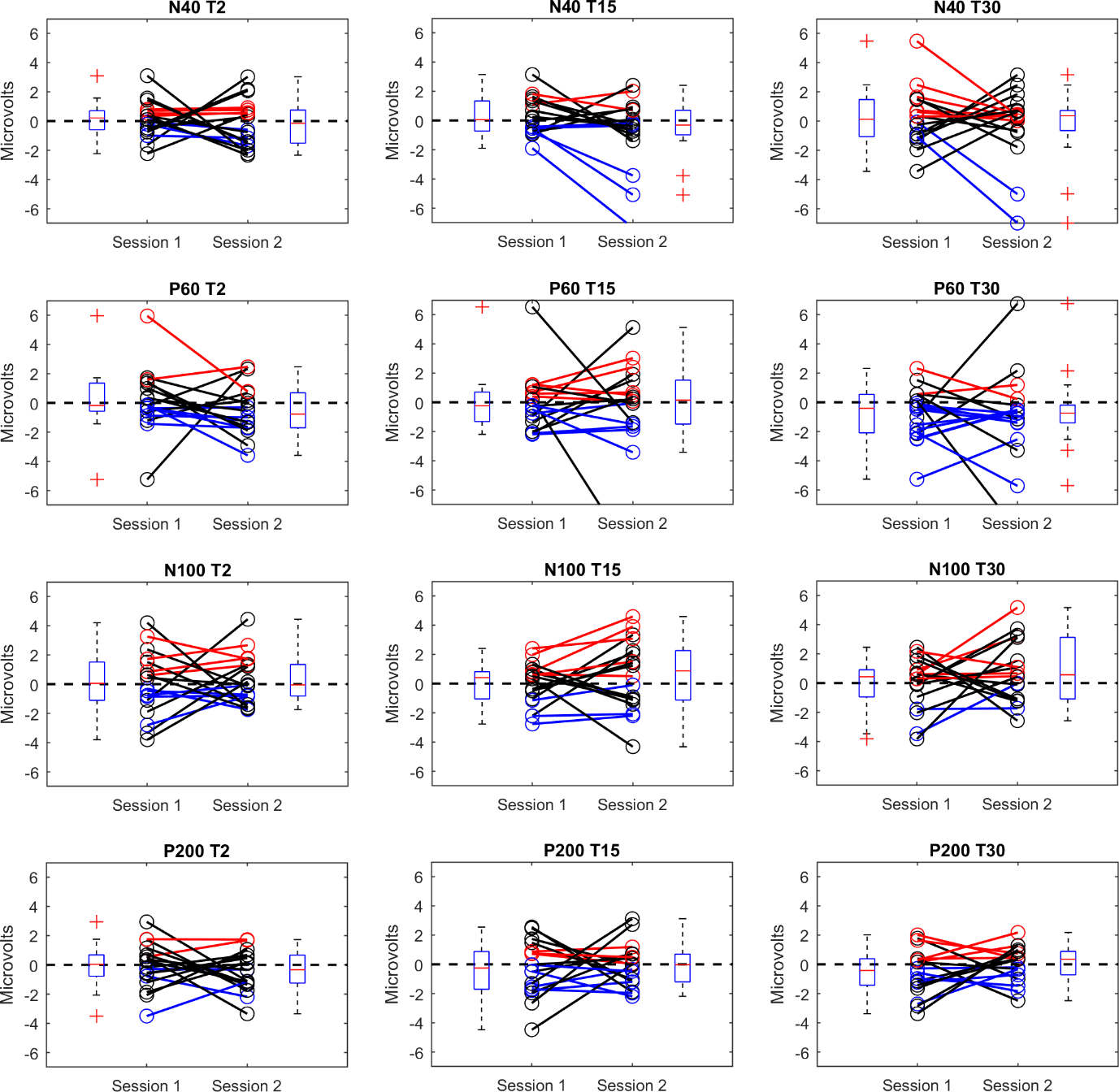

**Figure S11. Using ‘individual’ latencies**

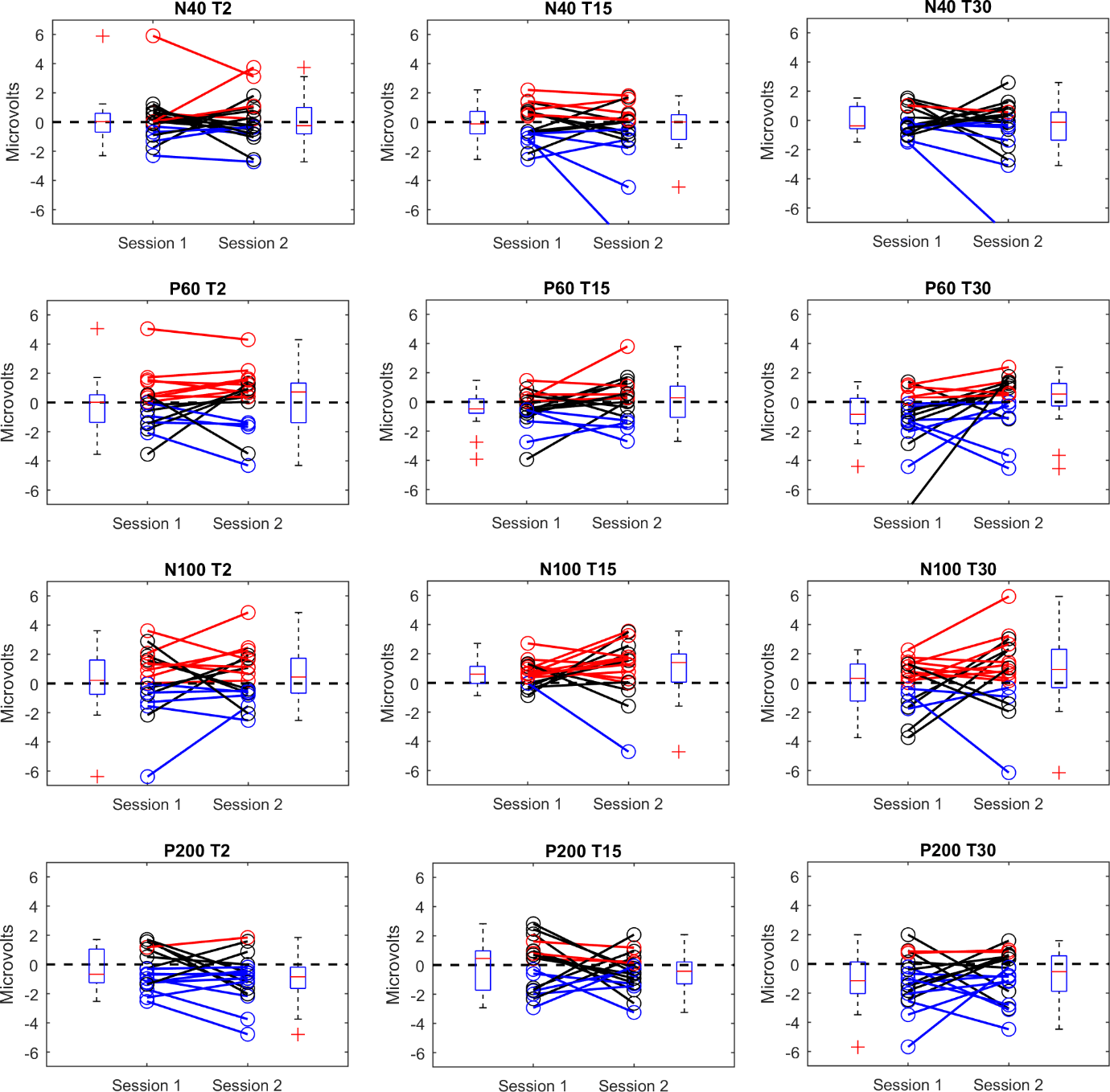

SHROUT, P. E. 1998. Measurement reliability and agreement in psychiatry. *Stat Methods Med Res,* 7**,** 301-17.
